# Surface Sensing Stimulates Cellular Differentiation in *Caulobacter crescentus*

**DOI:** 10.1101/844324

**Authors:** Rhett A. Snyder, Courtney K. Ellison, Geoffrey B. Severin, Christopher M. Waters, Yves V. Brun

**Affiliations:** Department of Biology, Indiana University, 1001 E. 3^rd^ Street, Bloomington, IN; Johns Hopkins School of Medicine Medical Science Training Program, Johns Hopkins University, Baltimore, MD; Lewis-Sigler Institute for Integrative Genomics, Princeton University, Princeton, NJ; Department of Biochemistry and Molecular Biology, Michigan State University, East Lansing, MI, USA; Department of Microbiology and Molecular Genetics, Michigan State University, East Lansing, MI, USA; Département de microbiologie, infectiologie et immunologie, Université de Montréal, C.P. 6128, succ. Centre-ville, Montréal (Québec) H3C 3J7

## Abstract

Cellular differentiation is a fundamental strategy used by cells to generate specialized functions at specific stages of development. The bacterium *C. crescentus* employs a specialized dimorphic life cycle consisting of two differentiated cell types. How environmental cues, including mechanical inputs such as contact with a surface, regulate this cell cycle remain unclear. Here, we find that surface sensing by the physical perturbation of retracting extracellular pilus filaments accelerates cell cycle progression and cellular differentiation. We show that physical obstruction of dynamic pilus activity by chemical perturbation or by a mutation in the outer membrane pilus pore protein, CpaC, stimulates early initiation of chromosome replication. In addition, we find that surface contact stimulates cell cycle progression by demonstrating that surface-stimulated cells initiate early chromosome replication to the same extent as planktonic cells with obstructed pilus activity. Finally, we show that obstruction of pilus retraction stimulates the synthesis of the cell cycle regulator, cyclic diguanylate monophosphate (c-di-GMP) through changes in the activity and localization of two key regulatory histidine kinases that control cell fate and differentiation. Together, these results demonstrate that surface contact and mechanosensing by alterations in pilus activity stimulate *C. crescentus* to bypass its developmentally programmed temporal delay in cell differentiation to more quickly adapt to a surface-associated lifestyle.

**Significance:** Cells from all domains of life sense and respond to mechanical cues [1–3]. In eukaryotes, mechanical signals such as adhesion and surface stiffness are important for regulating fundamental processes including cell differentiation during embryonic development [4]. While mechanobiology is abundantly studied in eukaryotes, the role of mechanical influences on prokaryotic biology remains under-investigated. Here, we demonstrate that mechanosensing mediated through obstruction of the dynamic extension and retraction of <u>t</u>ight <u>ad</u>herence (tad) pili stimulates cell differentiation and cell cycle progression in the dimorphic α-proteobacterium *Caulobacter crescentus*. Our results demonstrate an important intersection between mechanical stimuli and the regulation of a fundamental aspect of cell biology.

## Introduction

In multicellular organisms, cellular differentiation is required for the formation of complex tissues and organs [5]. In unicellular organisms, the ability to coordinate and control specialized cell morphologies and functions is critical for niche survival in diverse environments [6]. *C. crescentus* exhibits a dimorphic life cycle where asymmetric division results in the production of a non-reproductive, motile swarmer cell and a reproductive, non-motile stalked cell. In addition to their distinct reproductive states, each of these cell types possesses different polar structures. The swarmer cell is equipped with a single flagellum and multiple type IVc <u>t</u>ight <u>ad</u>herence (tad) pili at the same pole that are lost upon cellular differentiation into the stalked cell. Tad pili are subsequently replaced with a holdfast adhesin that mediates irreversible surface attachment and a thin cell-envelope extension called the stalk [7, 8].

Distinguishing characteristics between swarmer and stalked cells are partly due the action of the master response regulator, CtrA [8]. In swarmer cells, CtrA is phosphorylated and binds strongly to chromosomal sites near the origin of replication, preventing the initiation of DNA replication and thus locking cells in a non-reproductive, arrested G1 phase. During differentiation from swarmer to stalked cell, CtrA is dephosphorylated and proteolytically cleaved to allow for entry into S-phase and subsequent chromosome replication [8].

Regulatory control over differentiation is mediated by oscillating levels of c-di-GMP, a ubiquitous secondary messenger molecule that coordinates bacterial behavior in diverse species [9]. Newborn swarmer cells have low concentrations of c-di-GMP that slowly increase as they age. Between 20-40 mins post-division, a maximal level of c-di-GMP is observed, coinciding with holdfast synthesis and the transition from the motile to the sessile state. At the same time, a high level of c-di-GMP stimulates the dephosphorylation and deactivation of CtrA, allowing for chromosome replication as the swarmer cell differentiates [8].

c-di-GMP levels are controlled by the activity of the two histidine kinases PleC and DivJ, which localize at the swarmer and stalked pole of predivisional cells, respectively, and which dictate the distinct fates of the two progeny cells [10]. Delocalization of PleC and localization of DivJ at the incipient stalked pole during cell differentiation mediate the activation of the diguanylate cyclase PleD by phosphorylation, resulting in an increase in c-di-GMP.

Although the signal transduction network governing the transition from swarmer to stalked cell has been well described, whether surface attachment impacts this process is not known. Here, we demonstrate that inhibition of dynamic pilus activity stimulates c-di-GMP to initiate stalked cell development. We show that physical obstruction of pilus retraction and surface contact stimulate the initiation of DNA replication. We show that a mutation in the outer membrane pilus pore protein CpaC that partially disrupts pilus retraction stimulates holdfast synthesis and initiation of DNA replication in a PilA-dependent fashion. Finally, we show that physical obstruction of pilus retraction directly stimulates c-di-GMP synthesis by accelerating the delocalization of PleC and localization of DivJ at the incipient stalked pole, key steps in the activation of PleD and the production of c-di-GMP. Thus, by stimulating the synthesis of the holdfast [11] and cell differentiation, surface contact ensures that the permanently attached cell enters the stalked phase, which is best adapted for nutrient uptake on a surface [12].

## Results

### Obstruction of pilus retraction stimulates DNA replication initiation

Whether mechanical inputs can stimulate *C. crescentus* cell differentiation is unknown. Previous work has demonstrated that *C. crescentus* swarmer cells produce holdfast in response to surface contact independent of cell age [11, 13, 14], and recent findings suggests that this surface-stimulated holdfast synthesis is mediated by changes in type IVc tad pili dynamic activity upon binding of pili to a surface [11]. *C. crescentus* tad pili exhibit dynamic cycles of extension and retraction by polymerization and depolymerization of the major pilin subunit, PilA. Visualization of pili and their dynamic activity is achieved through knock-in cysteine mutation in PilA (Pil-cys) followed by the addition of thiol-reactive maleimide conjugates [11, 15, 16]. Dynamic activity of pilus fibers can be obstructed by the addition of bulky maleimide conjugates like polyethylene glycol maleimide (PEG-mal) to Pil-cys strains. In *C. crescentus*, obstruction of dynamic pilus activity through this method stimulates holdfast synthesis in the absence of surface contact, suggesting that the tension on surface-bound, retracting tad pili stimulates bacterial mechanosensing [11].

Because cell-cycle progression and cellular differentiation is concomitant with holdfast synthesis in planktonic cells, we hypothesized that surface contact may also accelerate the *C. crescentus* life cycle. A key marker for cell cycle progression is the initiation of DNA replication. We reasoned that should surface sensing stimulate initiation of DNA replication, cells with obstructed pili dynamics would have a higher DNA content compared to non-stimulated cells. To test this hypothesis, we incubated wild-type (WT) or Pil-cys cells with or without PEG-mal followed by rifampicin treatment to prevent new initiation of DNA replication while allowing for the completion of rounds of DNA replication that had already been initiated. We then labeled genomic DNA of treated cell cultures using SYTOX DNA-intercalating fluorescent dye and performed flow cytometry to quantify the DNA content of populations of cells. Swarmer cells arrested in the G1 phase harbor a single chromosome (1N), whereas cells that initiate chromosome replication prior to rifampicin treatment possess two chromosomes (2N). WT populations and untreated populations of the Pil-cys strain exhibited a ∼2-fold ratio of 2N:1N chromosome content. In contrast, the Pil-cys population treated with PEG-mal exhibited a ∼3-fold ratio of 2N:1N chromosome content (Figure 1A and B). Importantly, Pil-cys cells treated with polyethylene glycol lacking the thiol-reactive maleimide group (PEG) exhibited a ratio of 2N:1N genomic content similar to WT cells and untreated pil-cys cells. These results suggest that obstruction of pili dynamics stimulates the initiation of DNA replication.

**Figure 1.**
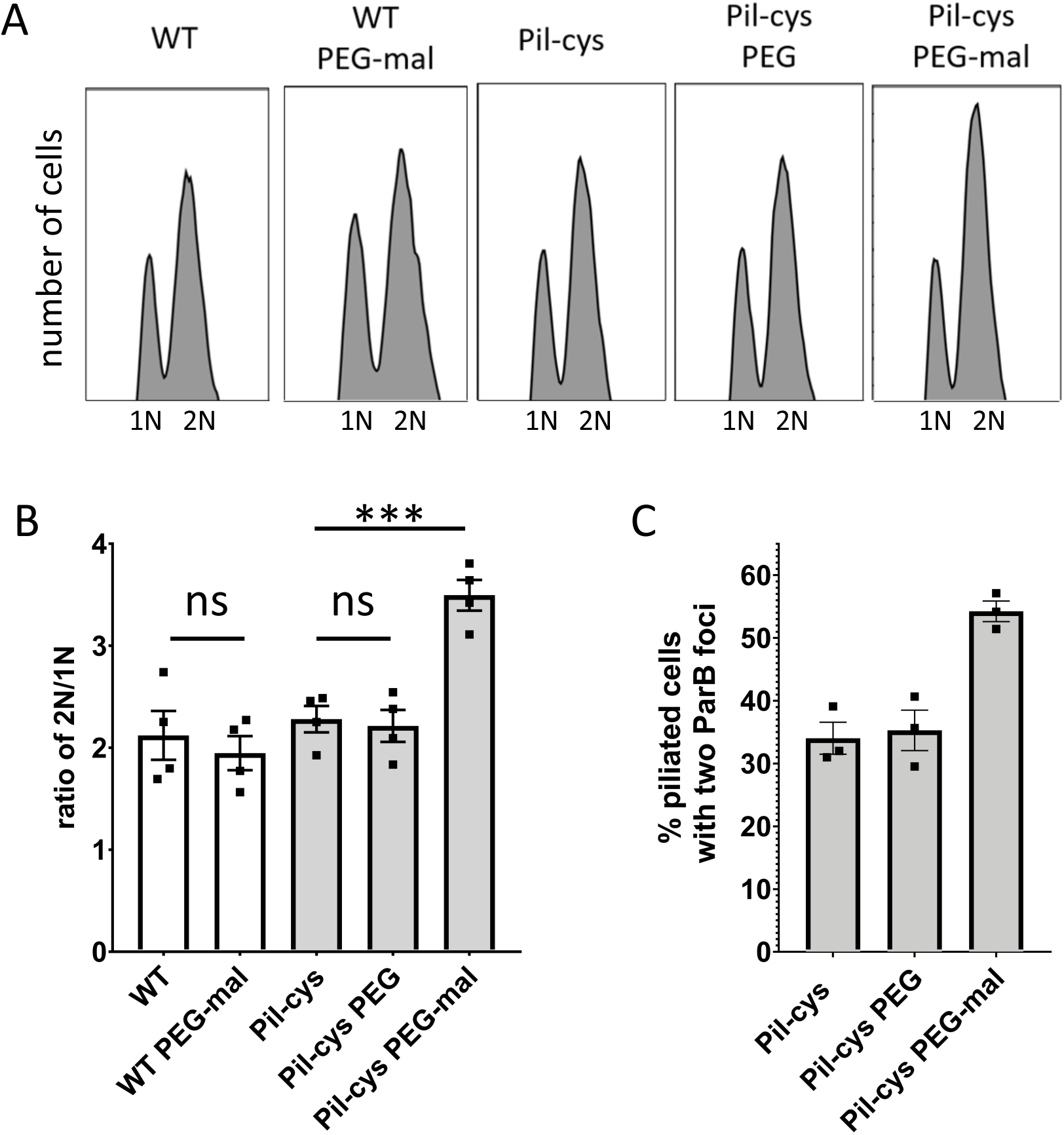
Obstruction of pilus retraction stimulates DNA replication initiation. (A) Representative flow cytometry plots showing chromosome content of cells quantified in (B). (B) Ratio of cells with two chromosomes (2N) to cells with one chromosome (1N) determined by flow cytometry analysis of genomic content. Bar graph shows the mean ± SEM of three independent, biological replicates. (C) Quantification of the percent of piliated cells with two ParB-mCherry foci. Bar graph shows the mean ± SEM of three independent, biological replicates. A minimum of 100 cells was quantified for each replicate. Statistical comparisons were made using Sidak’s multiple comparisons test. WT = wild type. ***P < 0.001, ns = not significant.

To confirm the above results, we tracked chromosome replication at the single-cell level. During S phase, the chromosomal <u>par</u>titioning system *parABS* in *C. crescentus* is involved in chromosome segregation. ParB dimers bind *parS* sequences adjacent to the origin of replication and subsequent interactions with cytoplasmic ParA helps to physically migrate the ParB-*parS*-DNA complex across the length of the cell [17]. To determine whether obstruction of pilus dynamics stimulates the initiation of DNA replication, we tracked the localization of the ParB in cells obstructed for pilus retraction with PEG-mal. For cells in G1 phase, a single ParB focus is observed at the flagellar pole where the origin of replication is localized. After initiation of DNA replication, a second ParB focus appears as newly-synthesized *parS* sites are bound by ParB dimers and translocated to the opposite cell pole. We thus examined the percentage of piliated cell with two ParB foci as a marker for cells that had initiated DNA replication. When treated with PEG-mal, the Pil-cys strain exhibited a 20% increase in the number of piliated cells with two ParB foci as compared to untreated and PEG-treated cells (Figure 1C). Taken together, these results suggest that obstruction of pili dynamics stimulates entry into the cell cycle.

### A mutation in the outer membrane pilus secretin that disrupts pilus retraction stimulates holdfast synthesis and initiation of DNA replication

Because chemical obstruction of pilus retraction through the addition of PEG-mal stimulates initiation of DNA replication, we reasoned that some mutants genetically deficient in pilus retraction would exhibit a similar phenotype. Since pili are terminally retracted prior to cellular differentiation, we hypothesized that stalked cells of a retraction mutant would exhibit an increase in the number of cells with pili localized at the tips of stalks where the outer membrane secretin CpaC remains after stalk synthesis [18]. Because retraction mutants in several species are hyperpiliated and hyperpilation results in increased surface attachment, we performed an unbiased genetic screen to enrich for mutants that attach more efficiently to surfaces. We then screened the enriched cell population for changes in pilus-dependent ϕCbK phage sensitivity because we assumed that a mutant deficient in pilus dynamics would be more resistant to pilus-dependent phage infection. From this screen, we isolated a mutant that harbored pili at the tips of stalked cells, indicative of obstructed pilus retraction and a failure to terminally retract its pili prior to cellular differentiation (Figure 2A and B). Whole genome sequencing revealed a mutation that mapped to the outer membrane pilus secretin gene, *cpaC*^G324D^.

**Figure 2.**
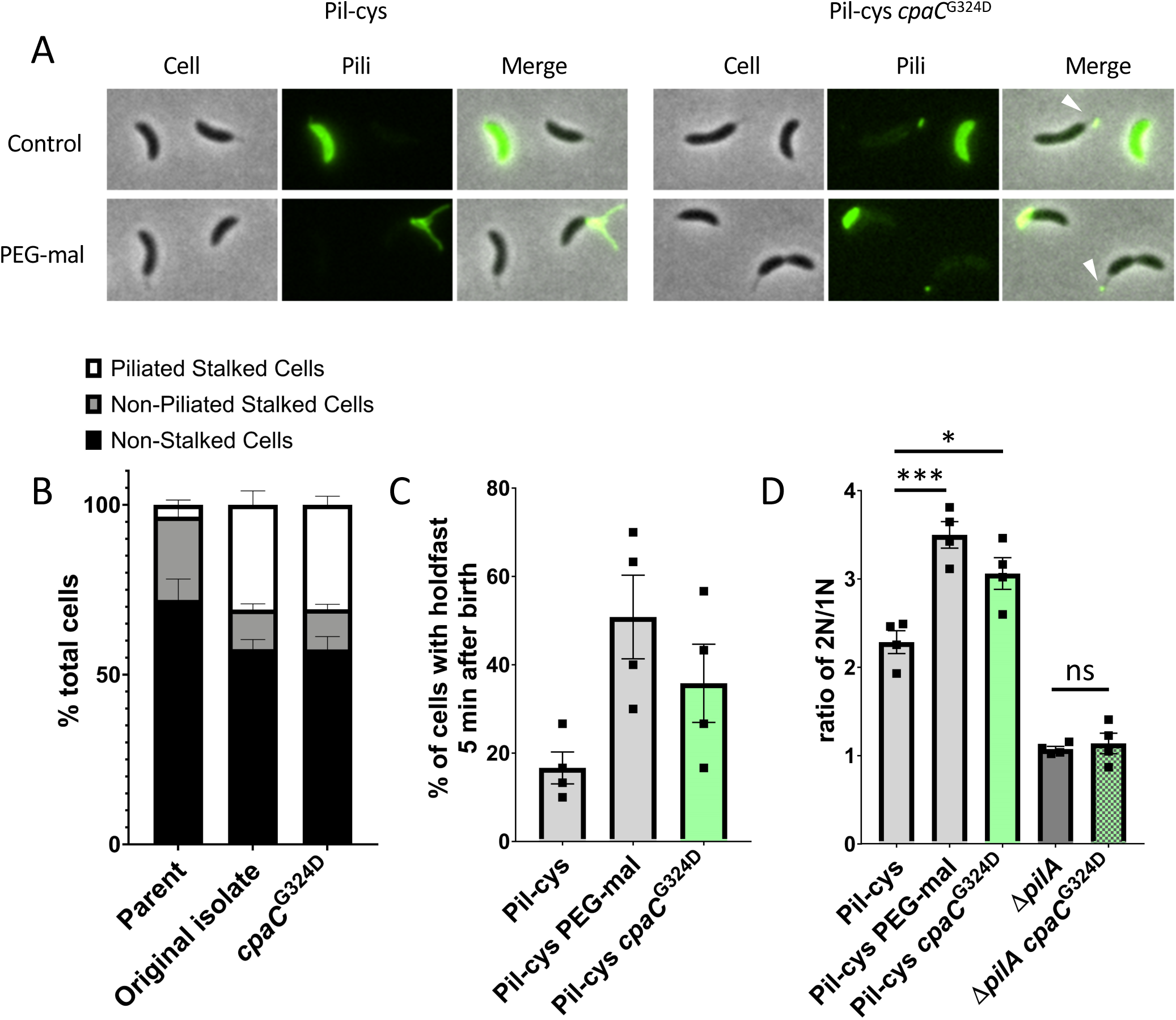
A mutation in the CpaC outer membrane pilus secretin partially obstructs pilus retraction and stimulates cell cycle progression and cellular differentiation. (A) Representative images of Pil-cys parent and strain containing *cpaC*^G324D^ mutation. Arrows indicate stalks with labeled pilus fibers attached to them. (B) Quantification of piliated stalk phenotype shown in (A). Data are from four independent, biological replicates and bar graphs show mean ± SEM. (C) Percent of synchronized cells that have made a holdfast by the start of the imaging experiment five min after birth. Bar graphs show mean ± SEM. Data are from four independent, biological replicates (n = 30 cells per replicate). (D) Ratio of cells with two chromosomes (2N) to cells with one chromosome (1N) determined by flow cytometry analysis of genomic content. Bar graph shows the mean ± SEM of four independent, biological replicates. Statistical comparisons were made using Sidak’s multiple comparisons test. *P < 0.05, ***P < 0.001, ****P < 0.0001.

To test whether the obstruction of pilus retraction mediated by the *cpaC*^G324D^ mutation stimulates cell cycle progression similarly to physical obstruction by PEG-mal treatment, we first quantified holdfast synthesis in mutant populations. In the *cpaC*^G324D^ mutant, approximately 36% of synchronized cells produced a holdfast within five minutes of birth as compared to 17% in cells with the wild-type allele of *cpaC* (Figure 2C). By comparison, 51% of cells obstructed for pilus retraction by the addition of PEG-mal synthesize a holdfast within five minutes of birth. These results suggest that the *cpaC*^G324D^ mutant is partially stimulated for surface sensing. Interestingly, the *cpaC*^G324D^ mutant appears only partially obstructed for pilus retraction as evidenced by fluorescent cell bodies (Figure 2A). Indeed, we have previously shown that cell body fluorescence in pil-cys cells labeled with fluorescent maleimide is dependent upon pilus retraction and dispersal of labeled pilins into an inner membrane pilin pool [11]. As the *cpaC*^G324D^ mutant exhibits both cell body fluorescence as well as pili at the tips of stalks, we infer that it is only partially obstructed for pilus retraction.

To test whether the *cpaC*^G324D^ mutant had an increase in DNA replication initiation similar to cells physically obstructed for pilus retraction, we measured the DNA content of *cpaC*^G324D^ mutants. We found that the *cpaC*^G324D^ mutant had an intermediate increase in the number of cells harboring two chromosomes compared to the PEG-mal treated pil-cys strain and WT, indicative of accelerated cell-cycle progression (Figure 2D). Importantly, a *cpaC*^G324D^ *pilA* double mutant lacking the major pilin subunit exhibited the same phenotype as a *pilA* mutant alone, demonstrating a dependence of cell-cycle acceleration of the *cpaC*^G324D^ mutant on the presence of PilA. These results suggest that obstruction of pili dynamics by the *cpaC*^G324D^ mutation stimulates both holdfast synthesis and entry into the cell cycle.

### Surface contact stimulates cell cycle progression

While physical obstruction of pilus retraction with PEG-mal or by *cpaC* mutation is inferred to simulate surface sensing in the absence of a surface, we sought to directly test whether surface contact stimulates cell cycle entry. Because cultures of *C. crescentus* harbor a mixture of undifferentiated swarmer cells, stalked cells, and predivisional cells at various stages of replication, we synchronized cultures of cells using a plate synchrony method to isolate newborn swarmer cells. We then tracked the timing of ParB duplication in surface-attached, planktonic, and PEG-mal treated planktonic populations (Figure 3A). Attached cells and planktonic cells treated with PEG-mal displayed similar ParB duplication times of 17 and 16.4 min after birth respectively, while untreated planktonic cells displayed a delay in ParB duplication of 19.7 min after birth (Figure 3B and C). Notably, the *cpaC*^G324D^ mutant that is genetically obstructed for pilus retraction exhibited ParB duplication at 17.6 minutes after birth, similar to both attached and PEG-mal-treated cells.

**Figure 3.**
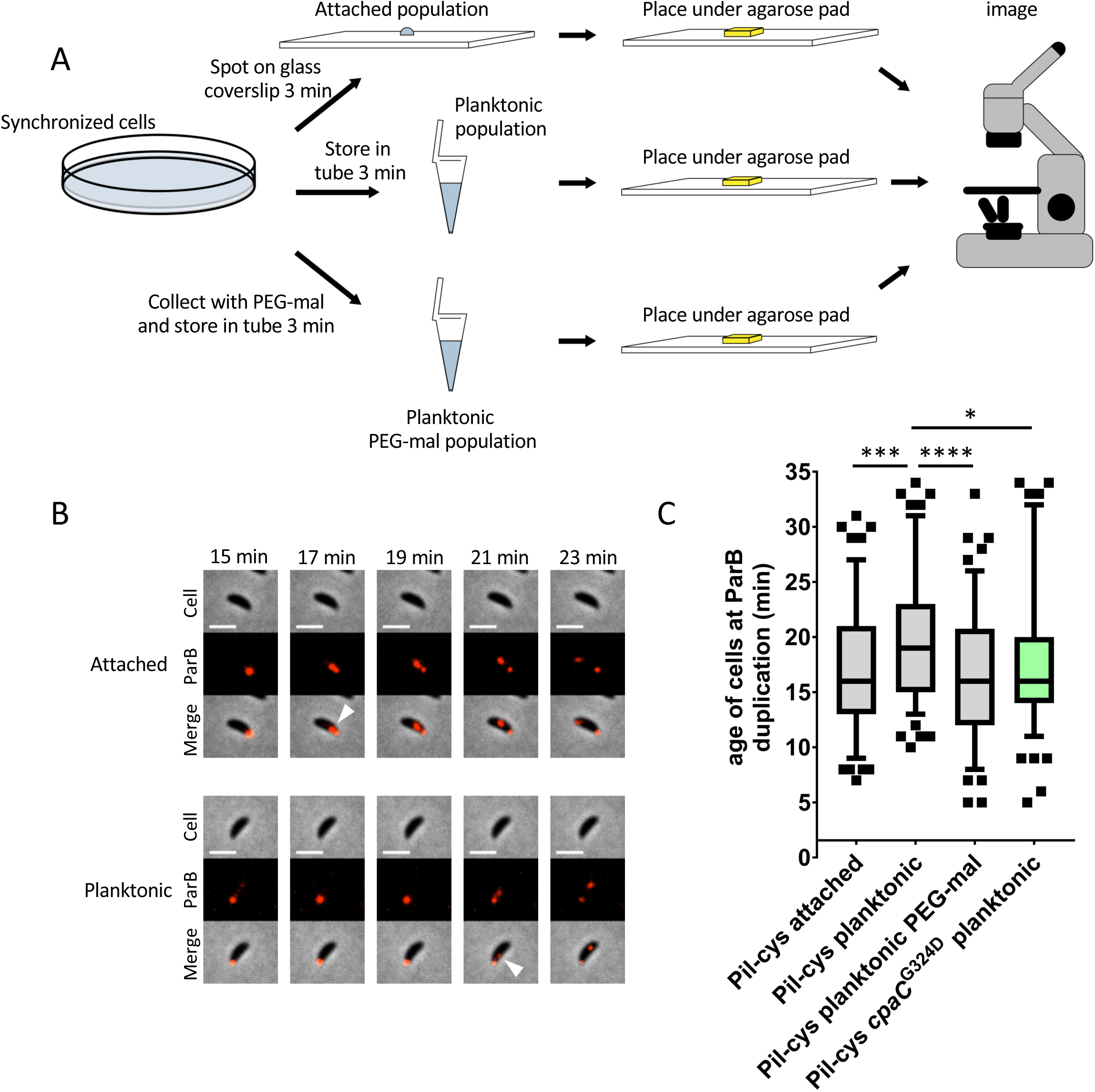
Surface contact stimulates cell cycle progression. (A) Schematic of experimental setup. (B) Representative time-lapse images of data shown in (C). Scale bars are 2 μm. White arrows indicate ParB duplication event. (C) Box and whisker plots show 5-95% confidence interval. Data are compiled from four independent, biological replicates (n = 30 cells per replicate). Statistical comparisons were made using Sidak’s multiple comparisons test. *P < 0.05, ***P < 0.001, ****P < 0.0001.

Taken together, our results indicate that swarmer cells that contact a surface, planktonic swarmer cells physically obstructed for pilus retraction, and planktonic swarmer cells with a mutation that obstructs pilus retraction differentiate ∼15% earlier than planktonic swarmer cells. We next sought to determine the mechanism by which obstruction of pili dynamics stimulates entry into the cell cycle.

### Obstruction of pilus retraction stimulates c-di-GMP synthesis by altering the activity of developmental regulators

The initiation of DNA replication and polar differentiation are tightly coupled during swarmer cell differentiation. This coupling is mediated in part by the histidine kinases PleC and DivJ, which antagonistically regulate the phosphorylation state of the single domain response regulator DivK in order to control entry into the cell cycle and the phosphorylation of PleD to stimulate c-di-GMP synthesis [19]. It was previously demonstrated that PleD is important for surface contact stimulation of holdfast synthesis [11], suggesting an increase of c-di-GMP upon surface sensing. We thus measured c-di-GMP concentrations of cell populations after obstruction of pilus activity (Figure 4A). WT cells lacking the Pil-cys mutation were unaffected by PEG-mal treatment while the Pil-cys strain exhibited a 50% increase in c-di-GMP concentration upon obstruction of pilus retraction. These results suggest that surface sensing by obstruction of pilus retraction is sufficient to stimulate the production of c-di-GMP.

**Figure 4.**
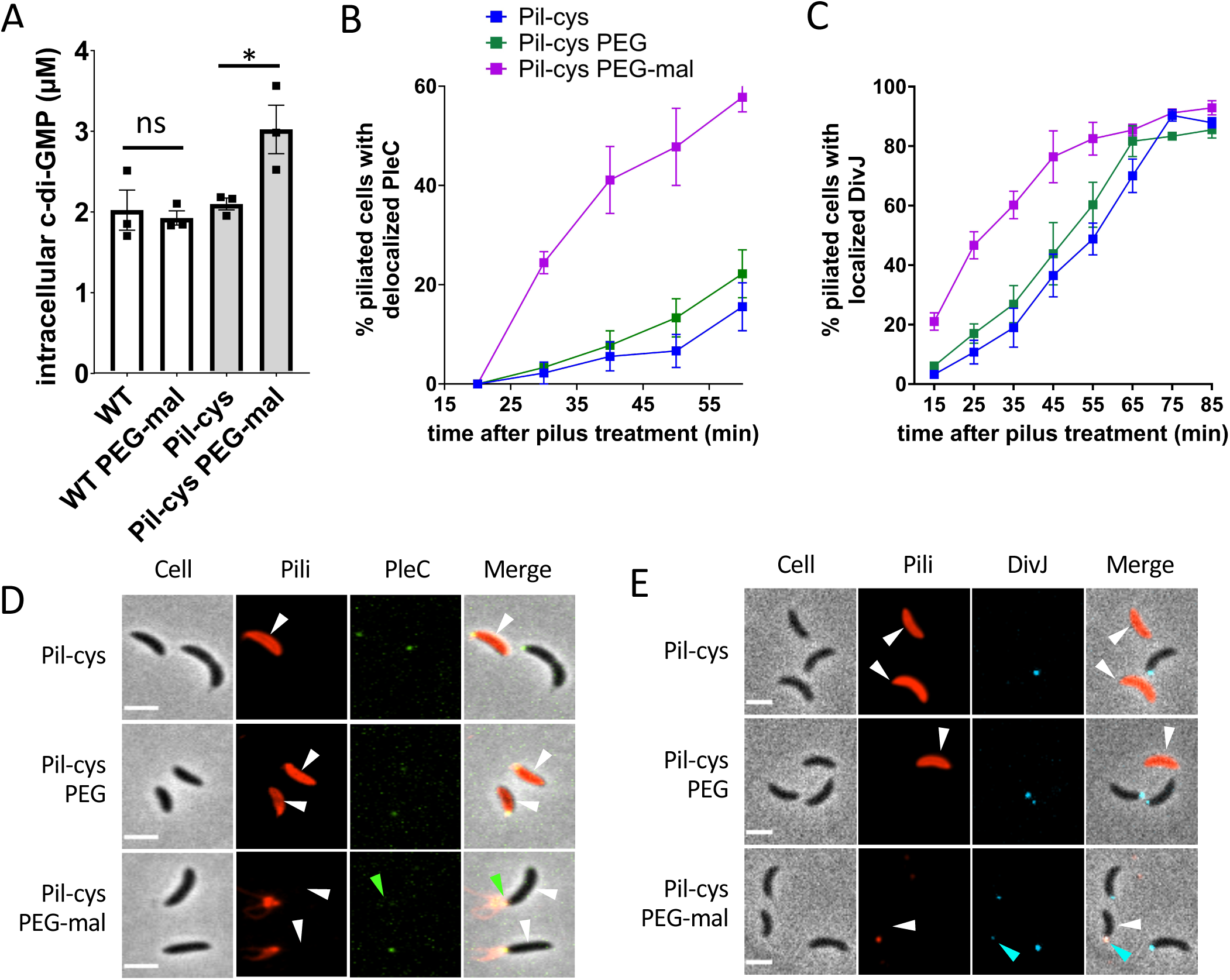
Obstruction of pilus retraction stimulates c-di-GMP synthesis by altering activity of developmental regulators. (A) Quantification of intracellular c-di-GMP concentrations of wild type and Pil-cys strains with PEG-mal treatment. Bar graph shows the mean ± SEM of three independent, biological replicates. Statistical comparisons were made using Sidak’s multiple comparisons test. WT = wild type. *P < 0.05, ns = not significant. (B) Percent of piliated cells with localized PleC at each time point. Error bars indicate mean ± SEM of three independent, biological replicates (n = at least 30 cells per replicate per time point). (C) Percent of piliated cells with localized DivJ at each time point. Error bars indicate mean ± SEM of four independent, biological replicates (n = at least 30 cells per replicate per time point). (D) Representative microscopy images of cells from data shown in (B). Green arrow represents delocalized PleC at the piliated pole. (E) Representative microscopy images of cells at the 25 min time point of data shown in (C). Blocked pili in Pil-cys PEG-mal treated samples appear as puncta due to shearing of filaments. Blue arrow indicates DivJ localization in piliated cell. White arrows indicate piliated cells. Scale bars are 2 μm.

c-di-GMP synthesis by PleD is spatially and temporally controlled by PleC and DivJ. Conveniently, the subcellular localization of PleC and DivJ correlates with their activity [8][19]. PleC is localized at the flagellar pole in swarmer cells where it acts as a phosphatase for DivK and PleD. PleC delocalizes and switches to a kinase during differentiation from the swarmer to stalked cell. During differentiation, DivJ interacts with its localization and activation factor SpmX to localize to the incipient stalked pole, where it phosphorylates DivK and PleD to start the cell cycle and stimulate cell differentiation [8]. To determine if surface sensing regulates this key regulatory switch, we tracked changes in PleC and DivJ localization in cells with obstructed pilus retraction as a proxy for their activity (Figure 4). Strains containing PleC-YFP or DivJ-CFP were treated with maleimide dye with or without PEG-mal, and piliated cells were tracked for changes in protein localization over time. The PEG-treated control and untreated Pil-cys populations exhibited a 20% increase in the number of cells with delocalized PleC by 60 min, while ∼60% of cells treated with PEG-mal delocalized PleC by the same time, demonstrating an acceleration in the PleC switch from phosphatase to kinase activity upon disruption of pilus dynamics (Figure 4B and D). DivJ also localized to the incipient stalked pole up to 20 min earlier in PEG-mal-treated samples in comparison to untreated or PEG-treated cells, showing that DivJ kinase activation is triggered by obstruction of pilus retraction (Figure 4C and E). These results demonstrate that the PleC-DivJ cell differentiation switch is stimulated by the obstruction of pilus retraction in addition to DNA replication and holdfast synthesis. Thus, mechanical cues upon surface contact stimulate differentiation of swarmer cells into stalked cells, which are better adapted for nutrient uptake on a surface [12].

## Discussion

It is becoming clear that mechanical signals from the environment have substantial impact on cell biology [20–22]. The mechanobiology of bacteria is an emerging field where we know little about the processes that can be modulated by mechanical signals and about how the signals are sensed and transduced. Here, we demonstrate that the mechanical cue of surface contact stimulates bacterial cell cycle progression and cell differentiation. We show that perturbation of pilus dynamic activity through surface contact, physical obstruction, or mutation of the pilus outer membrane pore stimulates DNA replication initiation and that physical obstruction of pilus dynamics causes a spike in c-di-GMP synthesis. These results suggest that surface contact causes an increase of c-di-GMP as a consequence of perturbation of pili dynamics and that this increase in c-di-GMP stimulates cell cycle progression and cell differentiation. It was previously shown that PleD is the main diguanylate cyclase responsible for the increase of c-di-GMP during swarmer to stalked cell differentiation [19]. C-di-GMP production by PleD stimulates the ShkA-TacA phosphorylation cascade, ultimately creating a positive feedback loop that results in increased PleD activity and ensures irreversible commitment to cell differentiation [23]. The activity of PleD is modulated by the histidine kinases PleC and DivJ, whereby PleC dephosphorylates PleD to inhibit its activity and DivJ phosphorylates PleD to activate it [19]. Delocalization of PleC and localization of DivJ at the incipient stalked pole result in an increase in PleD phosphorylation and c-di-GMP level, which triggers cell differentiation. PleC and DivJ similarly antagonistically modulate the phosphorylation state of the single domain response regulator DivK to ultimately control CtrA activity and chromosome replication [19]. We show that obstruction of pilus dynamics accelerates delocalization of PleC and localization of DivJ at the incipient stalked pole, which is expected to increase c-di-GMP and thereby stimulate entry into the cell cycle and cell differentiation (Figure 5). At the same time, surface contact also stimulates holdfast synthesis through flagellum motor interference and obstruction of pili dynamics, causing a spike in c-di-GMP that allosterically activates the holdfast polysaccharide glycosyltransferase HfsJ to stimulate holdfast synthesis [11, 13, 14].

**Figure 5.**
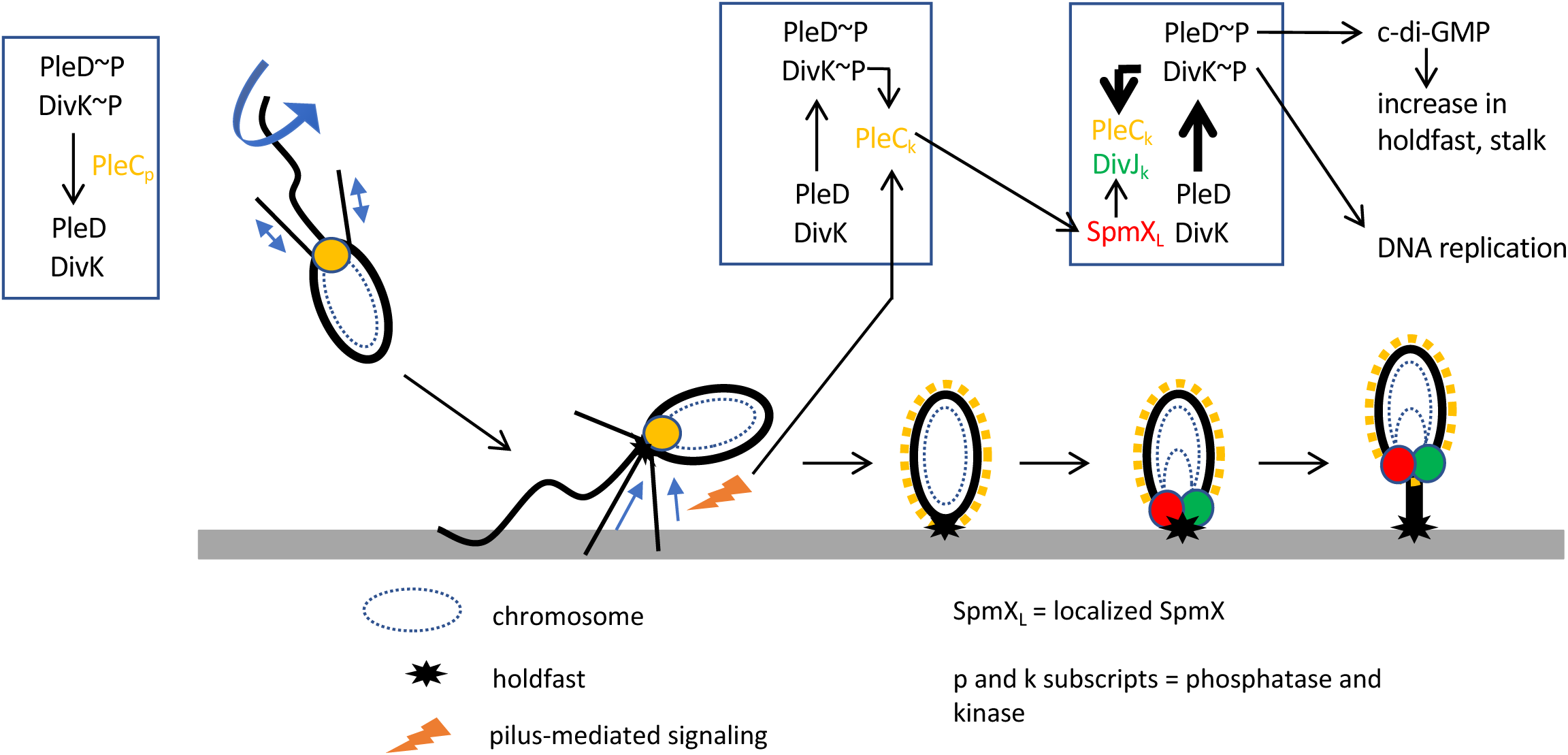
Model of cell cycle acceleration upon surface contact. Surface sensing through alterations in pilus retraction upon surface binding stimulates the PleC switch from phosphatase to kinase activities of PleC which occurs upon PleC delocalization. This in turn stimulates the localization of SpmX which recruits the kinase DivJ. DivJ phosphorylates PleD and DivK, resulting in the production of in c-di-GMP and the stimulation of DNA replication respectively. Increased c-di-GMP production from phosphorylated PleD results in more holdfast synthesis and stalk growth.

When newborn swarmer cells swim to a surface, the DivJ kinase is not yet localized nor active [24] and PleC is localized at the pole bearing pili and the flagellum where it acts as a phosphatase to dephosphorylate PleD, preventing its activation and localization [19]. The accelerated delocalization of PleC and its concomitant switch to a kinase is therefore likely to be the first step in the stimulation of PleD activity. Furthermore, the elevation of DivK∼P concentration stimulates DivJ kinase, causing a positive feedback loop between PleC and DivJ that leads to an increase in both DivK∼P and PleD∼P [19]. This synergy is also likely potentiated by PleC’s positive action on the localization and activation of DivJ by SpmX [24]. The co-localization of PleC with the pili and flagellum suggests that there may be crosstalk between the two surface contact sensory systems and PleC, providing an integration of holdfast synthesis, initiation of DNA replication, and cell differentiation upon surface contact. In support of this model, data from a parallel study by Del Medico et al. suggest that a PilA signal sequence is involved in stimulating c-di-GMP synthesis to trigger cell cycle progression through the PleC-PleD signaling cascade [25]. From an ecological perspective, accelerated cellular differentiation after surface contact and permanent attachment likely benefits *C. crescentus* by activating the pathway that stimulates stalk synthesis. Indeed, the synthesis of a thin stalk is thought to improve nutrient uptake capacity in the diffusion-limited environment of a surface [12, 26]. A recent study demonstrated that some bacteria can sense and respond to changes in liquid flow rates [21], and accelerated stalk synthesis may also provide an advantage to surface-associated cells by allowing better access to environmental flow conditions [27].

Finally, our results are an important milestone in understanding how cells sense and respond to their environments by highlighting that physical cues can influence the hardwired circuitry of cellular differentiation and reproduction. Elucidating how cells sense and transduce the inputs from mechanical stimuli will be critical for determining how mechanical stimuli influence intracellular processes.

## Materials and methods

### Bacterial strains, plasmids, and growth conditions

Bacterial strains and primers used in this study are listed in Supplemental Table 1. *C. crescentus* strains were grown at 30°C in peptone yeast extract (PYE) medium [28]. *Escherichia coli* DH5α (Bioline) were used for cloning and grown in lysogeny broth (LB) medium at 37°C supplemented with 25 μg/ml kanamycin when appropriate.

Plasmids were transferred to *C. crescentus* by electroporation, transduction with ΦCr30 phage lysates, or conjugation with S-17 *E. coli* strains as described previously [29]. In-frame allelic substitutions were made by double homologous recombination using pNPTS-derived plasmids as previously described [30]. Briefly, plasmids were introduced to *C. crescentus* and then two-step recombination was performed using sucrose and kanamycin resistance or sensitivity as a selection for each step. Mutants were verified through a combination of sequencing and microscopy phenotyping.

For construction of the pNPTS-derived plasmids, ∼500 bp flanking regions of DNA on either side of the desired mutations were amplified from *C. crescentus* genomic DNA. Point mutations were built into the UpR and DownF primers as indicated in Supplemental Table 1. Upstream regions were amplified using UpF and UpR primers while downstream regions were amplified using DownF and DownR primers. The resulting DNA was purified (Qiaquick, Zymo Research) and assembled in pNPTS138 that had been digested with restriction enzyme *Eco*RV (New England Biolabs) using HiFi Assembly Master Mix (New England Biolabs). For construction of pNPTS138*hfsA+*, the entire *hfsA* gene and ∼500 bp of both up and downstream flanking DNA was amplified from strain FC764 [31] and cloned into pNPTS138 as described above for use in restoring holdfast synthesis in NA1000 strains.

### Cyclic-di-GMP quantification

Cyclic di-GMP was quantified as described previously [32]. Briefly, strains were grown to early-log growth phase (OD_600_ 0.15-0.25) in PYE medium. One ml of culture was centrifuged for five min at 21,000 x *g* and the supernatant was removed. Cell pellets were resuspended in 200 μl cold extraction buffer (1:1:1 mix of methanol, acetonitrile, and distilled H_2_O + 0.1 M formic acid) and incubated at -20°C for 30 min. Samples were then centrifuged at 21,000 x *g* to pellet cell debris, and the supernatant was transferred to a fresh tube and stored at -80°C until use. Experimental extraction solutions were desiccated overnight in a SpeedVac, re-solubilized in 100 μl of ultrapure water, briefly vortexed and centrifuged for 5 min at 21,000 x *g* to pellet insoluble debris. The clarified extract solutions were transferred to sample vials and analyzed by UPLC-MS/MS in negative ion-mode electrospray ionization with multiple-reaction monitoring using an Acquity Ultra Performance LC system (Water Corp.) coupled with a Quattro Premier XE mass spectrometer (Water Corp.) over an Acquity UPLC BEH C18 Column, 130 angstrom, 1.7 μm, 2.1 mm x 50 mm. c-di-GMP was identified using precursor > product masses of 689.16 > 344.31 with a cone voltage of 50.0 V and collision energy of 34.0 V. Quantification of c-di-GMP in sample extracts was determined using a standard curve generated from chemically synthesized c-di-GMP (AXXORA). The standard curve solutions were prepared using twofold serial dilutions of c-di-GMP (1.25 μM – 19 nM) in ultra-pure water that were further diluted 1:10 into biological extracts (final c-di-GMP concentrations: 125 nM to 1.9 nM) from a low c-di-GMP strain of *C. crescentus* lacking several diguanylate cyclases (c-di-GMP0) described previously [9] which had been grown, harvested, extracted, desiccated and solubilized in ultrapure water in tandem with the experiments samples described above. General UPLC buffer preparations, chromatographic gradients and MS/MS parameters were performed using a previously published method [33]. Intracellular concentrations of c-di-GMP were calculated as described previously [34] assuming *C. crescentus* average cellular volume of 6.46 × 10^−16^ L. The total number of cells present in each extraction was calculated by normalizing OD_600_ for each sample to the average CFUs found for NA1000 cultures grown to an OD_600_ 0.2 (2 × 10^9^ CFU/ml).

### Quantification of piliated cells with two ParB foci

Bacterial cultures were grown to an OD_600_ of 0.2-0.4 and labeled for pili as described previously [11]. Briefly, 100 μl of cultures were labeled with 25 μg/ml AlexaFluor 488 C_5_ maleimide dye (AF488-mal)(Thermofisher) for five min at room temperature. To block pilus retraction, cells were incubated simultaneously with AF488-mal and 500 μM of methoxypolyethylene glycol maleimide (5000 Da)(PEG-mal)(Sigma) for five min at room temperature. Cells were centrifuged at 5,200 x *g* for one min, the supernatant was discarded, and the pellet was then washed with 100 μl of PYE and centrifuged again. The supernatant was removed, and the cells were concentrated in 5-8 μl of PYE. One μl of washed, labeled cells was spotted onto a 60 × 22 glass coverslip and imaged under a 1% agarose (SeaKem) pad made with sterile, distilled water before imaging. Imaging was performed on an inverted Nikon Ti-2 microscope using a Plan Apo 60X objective, a GFP/DsRed filter cube, a Hamamatsu ORCAFlash4.0 camera, and Nikon NIS Elements Imaging Software. Quantification of piliated cells and number of ParB foci was performed manually using NIS Elements Analysis software.

### Quantification of genomic content in populations of cells

Bacterial cultures were grown to an OD_600_ of 0.2-0.4 and were left untreated or were treated with either 500 μM of PEG5000-mal or 500 μM polyethylene glycol (∼5000 Da)(Sigma). After pilus treatment, cells were incubated with 15 μg/ml of rifampicin for 3 h to prevent new cycles of DNA initiation. 1.5 ml of culture was concentrated into 180 μl PBS (phosphate buffered saline) and fixed with 420 μl 100% ethanol at 4°C for one hour. After fixation, cells were centrifuged at 5,200 x *g* and washed once with 600 μl PBS. Cells were finally resuspended in 600 μl PBS and 2.5 μM SYTOX Green Nucleic Acid Stain (Thermofisher) was added preceding stationary incubation overnight at 4°C. Fluorescence intensity and light scattering were quantified by flow cytometry using the FACSCalibur at the IUB FACS facility and data were analyzed using FlowJo software.

### Quantification of PleC and DivJ localization patterns

Pili were labeled with AF594-mal (Thermofisher) and treated with either PEG-mal or PEG as described above. For tracking DivJ-CFP localization after pilus treatment, cells were placed in a static, 30°C incubator. Every 10 min, one μl of sample was spotted onto a glass coverslip and imaged using DsRed/CFP filter settings under 1% agarose pads made with sterile, distilled water as described above. For tracking PleC delocalization, cells were spotted onto a glass coverslip and placed under a 1% agarose pad made with PYE and an initial image was taken using DsRed filter settings to identify piliated cells. Cells were then imaged using YFP filter settings every two min to track PleC-YFP delocalization. Quantification of piliated cells with delocalized PleC or localized DivJ was performed manually using NIS Elements Analysis software.

### Identification of mutant deficient in pilus retraction

A subculture-based forward genetic screen was performed to enrich for mutants efficient in holdfast-independent surface attachment. Ten replicates of a parent Pil-cys strain lacking the holdfast-synthesizing genes (Δ*hfsDAB*) was grown in five ml of PYE in glass tubes to stationary phase. Cultures were then dumped and lightly washed with PYE to remove loosely bound cells. The tubes were then refilled with five ml of PYE and again grown to stationary phase, and this was repeated until turbid growth was observed after overnight growth (23 days). Cultures were then streaked out onto PYE agar plates to isolate individual mutants. Isolates were then tested for changes in phage sensitivity to the pilus-dependent phage ΦCbK, and those exhibiting an alteration from wildtype sensitivity were sequenced to identify mutations. Whole genome sequencing and mutant identification was performed as described previously [35] with the exception that sequencing reads were mapped to the genome of *C. crescentus* NA1000 (NC_011916.1).

### Phage sensitivity assays

Phage sensitivity assays were performed as described previously [36]. Briefly, five μl of ΦCbK phage dilutions were spotted onto lawns of growing *C. crescentus* strains. Lawns were made by adding 200 μl of stationary phase cultures to three ml of melted top agar (0.5% agar in PYE) and spread over 1.5% PYE agar plates. After the top agar solidified, five μl of phage dilutions in PYE were spotted on top. Plates were grown for two days at 30°C before imaging.

### Cell synchronization and surface stimulation experiments

Cells were synchronized as described previously [11] with some modifications. Briefly, 50 ml of PYE in a 15 cm polystyrene petri dish was inoculated with one ml of overnight culture of the indicated holdfast-synthesizing strain expressing ParB-mCherry and incubated for 16 h at room temperature at 70 rpm on an orbital shaker. Four hours prior to experiments, the petri dish was washed with 50 ml of sterile, distilled water. 50 ml of PYE medium was added to the petri dish and incubated at room temperature shaking for an additional four hours. Just before use, the petri plate was washed twice with 100 ml of distilled water, and then one ml of PYE (containing 500 μM PEG-mal where indicated) was added to the petri plate and harvested after one minute to collect newborn swarmer cells. For planktonic populations, the one ml of PYE containing newborn swarmer cells was added to a 1.7 ml centrifuge tube and left stationary at room temperature for three minutes before 1 μl was spotted onto a coverslip and imaged under a 1% agarose pad made with PYE and containing 0.5 μg/ml AF488-WGA to label holdfasts. For surface-attached cells, 1 μl of the harvested newborn swarmer cells was spotted onto a glass coverslip and left stationary at room temperature for three minutes before the addition of the 1% agarose pad. Agarose pads do not stimulate surface-contact responses as reported elsewhere [32], and we found that allowing cells to attach to the glass coverslip for three minutes before the addition of the pad was critical for observing a surface-stimulated response. Time-lapse images of ParB-mCherry foci and holdfasts labeled with AF488-WGA in the agarose pad were captured once per minute over 35 min using the same settings described above. Holdfast and ParB-mCherry duplication events were quantified manually using NIS Elements Analysis software.

## Acknowledgements

We thank A. Dalia and C. Berne and members of the Gitai lab for helpful discussions regarding the manuscript. We thank the Center for Genomics and Bioinformatics at Indiana University for whole genome sequencing and SNP mutant identification. We also thank the FACS Core Facility at IUB for training and assistance in flow cytometry experiments. We thank D. Kysela for construction of strain YB7341.

## Funding

This study was supported by grant R35GM122556 from the National Institutes of Health and by a Canada 150 Research Chair in Bacterial Cell Biology to YVB, by grant R01GM109259 from the National Institutes of Health to CMW, and by National Science Foundation fellowship 1342962 to CKE;

## Author contributions

RAS, CKE, GBS, CMW and YVB designed the experiments. RAS, CKE, and GBS performed the experiments. All authors analyzed the data. RAS, CKE, and YVB wrote the manuscript. All authors contributed to editing the manuscript;

## Competing interests

The authors declare no competing interests;

## Data and materials availability

All data is available in the main text or the supplementary materials.

---

**Supplemental Table 1.**
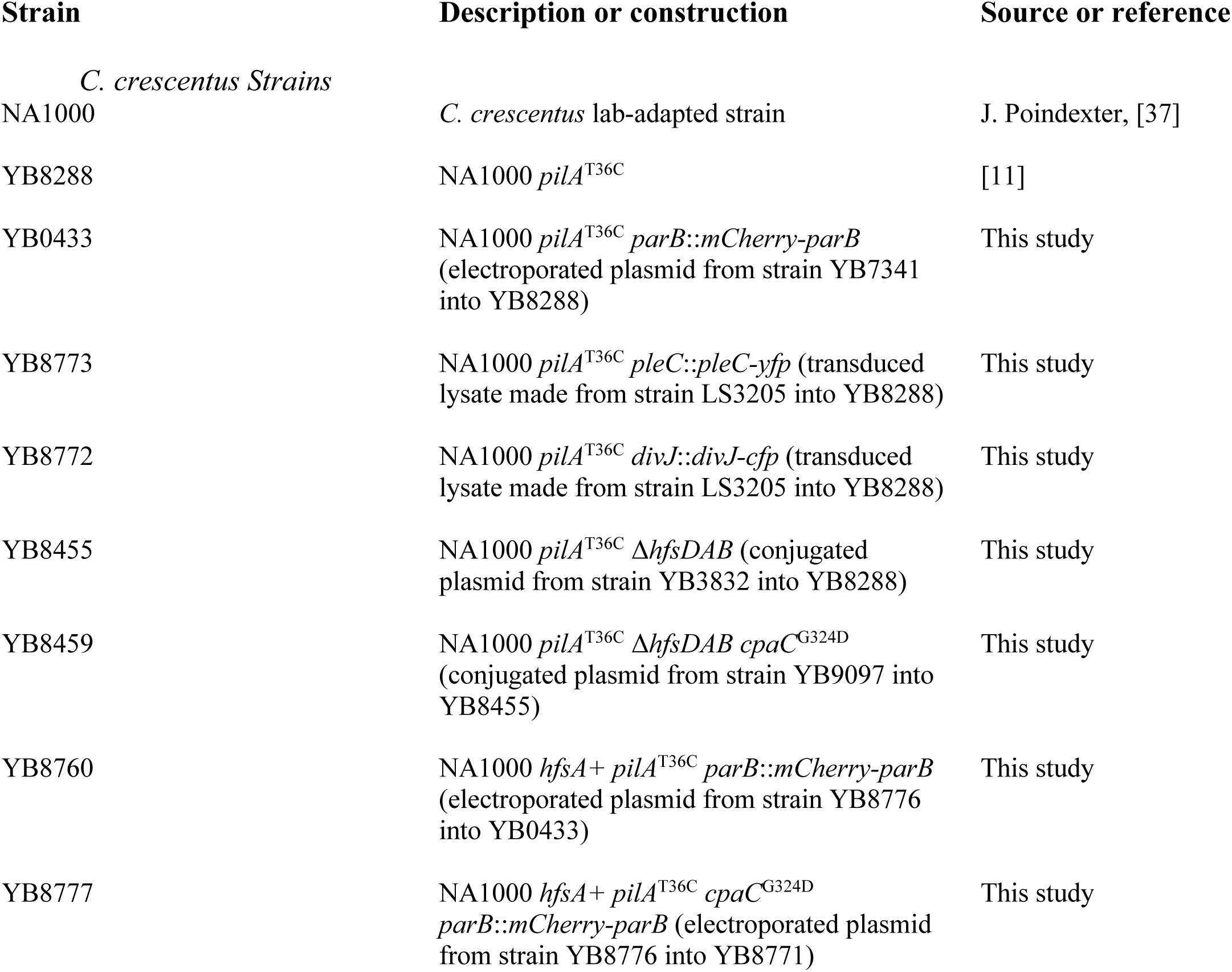
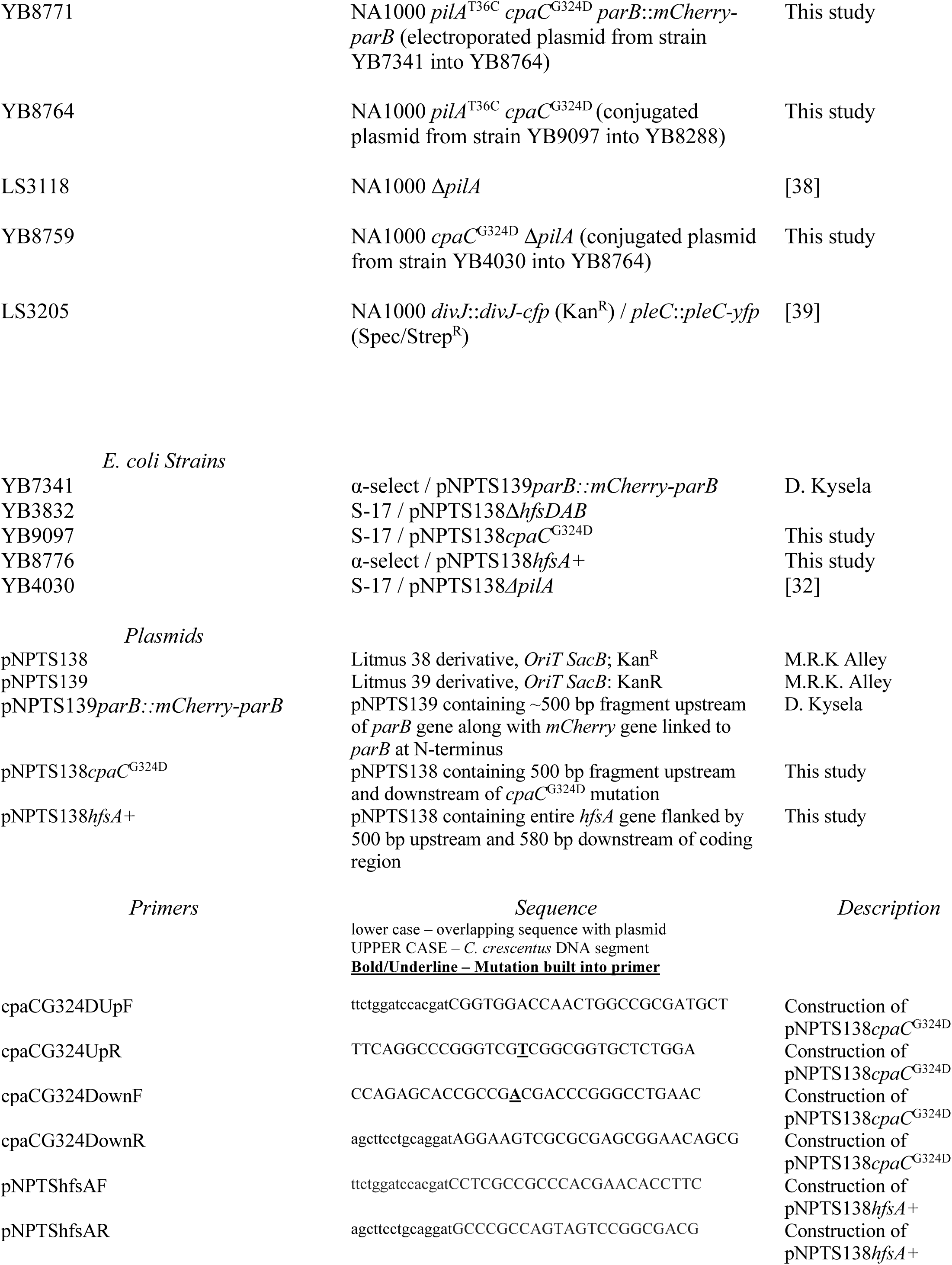
Strains, plasmids, and primers used in this study.

